# Considerations for furbearer trapping regulations to prevent grizzly bear toe amputation and injury

**DOI:** 10.1101/2021.07.06.450999

**Authors:** Clayton Lamb, Laura Smit, Bruce Mclellan, Lucas M. Vander Vennen, Michael Proctor

## Abstract

Science and adaptive management form crucial components of the North American model of wildlife management. Under this model, wildlife managers are encouraged to update management approaches when new information arises whose implementation could improve the stewardship and viability of wildlife populations and the well-being of animals. Here we detail a troubling observation of several grizzly bears with amputated toes in southeast British Columbia and assemble evidence to inform management strategies to remedy the issue. During the capture of 59 grizzly bears, we noticed that four individuals (~7%) had amputated toes on one of their front feet. The wounds were all healed and linear in nature. Further opportunistic record collection revealed that similar examples of amputated toes occurred beyond our study area, and that furbearer traps were frequently responsible for toe loss. We found evidence that seasonal overlap between the active season for grizzly bears and the fall trapping seasons for small furbearers with body grip traps and for wolves with foothold traps may explain the issue. Multiple options to reduce or eliminate this issue exist, but have varying degrees of expected efficacy and require differing levels of monitoring. The most certain approach to greatly reduce the issue is to delay the start of the marten trapping season until December 1, when most bears have denned, instead of opening the season on or prior to November 1, when more than 50% of bears are still active. Additionally, innovative solutions, such as narrowing trap entrances to exclude bear feet while still allowing entrance of target furbearers, have the potential to minimize accidental capture of bears, but the effectiveness of these approaches is unknown. Experimental evidence suggested that better anchoring traps was not a viable solution. Solutions that do not involve season changes will require monitoring of efficacy and compliance to ensure success.

## IDENTIFYING AN ISSUE

Grizzly bears (*Ursus arctos*) are wide-ranging mammals that are highly motivated to rapidly ingest high-energy foods in preparation for denning (McLellan 2011). In the Rocky Mountains of southern Canada, grizzly bears occupy large home ranges (200-1300 km^2^) where they search for mates and seasonal foods (Graham and Stenhouse 2014). Key natural foods for grizzly bears include high-calorie fruits, abundant herbaceous vegetation, ungulates (killed or scavenged), ants, and roots (McLellan and Hovey 1995, Munro et al. 2006). In addition to these natural foods, grizzly bears are well known to take advantage of poorly managed human foods such as residential fruit trees, garbage, roadkill, grain, and livestock (Craighead and Craighead 1972, Lamb et al. 2017, 2019, Morehouse et al. 2020). This food-motivated behaviour is generally adaptive and allows grizzly bears to increase their weight by about 35% in the 6-8 months of non-denning season (McLellan 2011), but bears can also be attracted to, and allowed to feed on, human-sourced foods which can create human and bear safety issues (Lamb et al. 2020).

The challenges to human-bear coexistence are exacerbated when high-density grizzly bear populations co-occur in human settled areas with abundant natural foods and accessible human-sourced foods. One area that fits this description well is the southeast corner of British Columbia (BC), Canada. This region is home to high-density grizzly bear populations, productive habitat, and growing rural and urban communities. In response to recent grizzly bear population declines, ecological trap dynamics, and high human-caused mortality rates in this area (Lamb et al. 2017, 2019, 2020), we initiated a radio collaring project to better understand sources of grizzly bear mortality and examine how animals’ use of the landscape influenced population dynamics. Between 2016 and 2020, we captured and radio collared 59 grizzly bears in the Elk Valley (near Fernie, BC, see Fig 1A). Captures were in accordance with University of Alberta Animal Ethics #AUP00002181 and Province of British Columbia Capture Permit #CB17-264200. Here we focus on a particular observation from this live capture work that has important ramifications: several of the bears had amputated toes. That is the toes appeared to have been cut off and then healed. This observation is important because ethics and values have a major role in wildlife management in British Columbia and resulted in closing a decades-long sustainable grizzly bear hunt in 2018, despite a large and viable population (Auditor General 2017, McLellan et al. 2017, Hatter et al. 2018). Identifying how these toes were amputated and mitigating the source of amputation became one of the objectives of our study.

**Figure 1.**
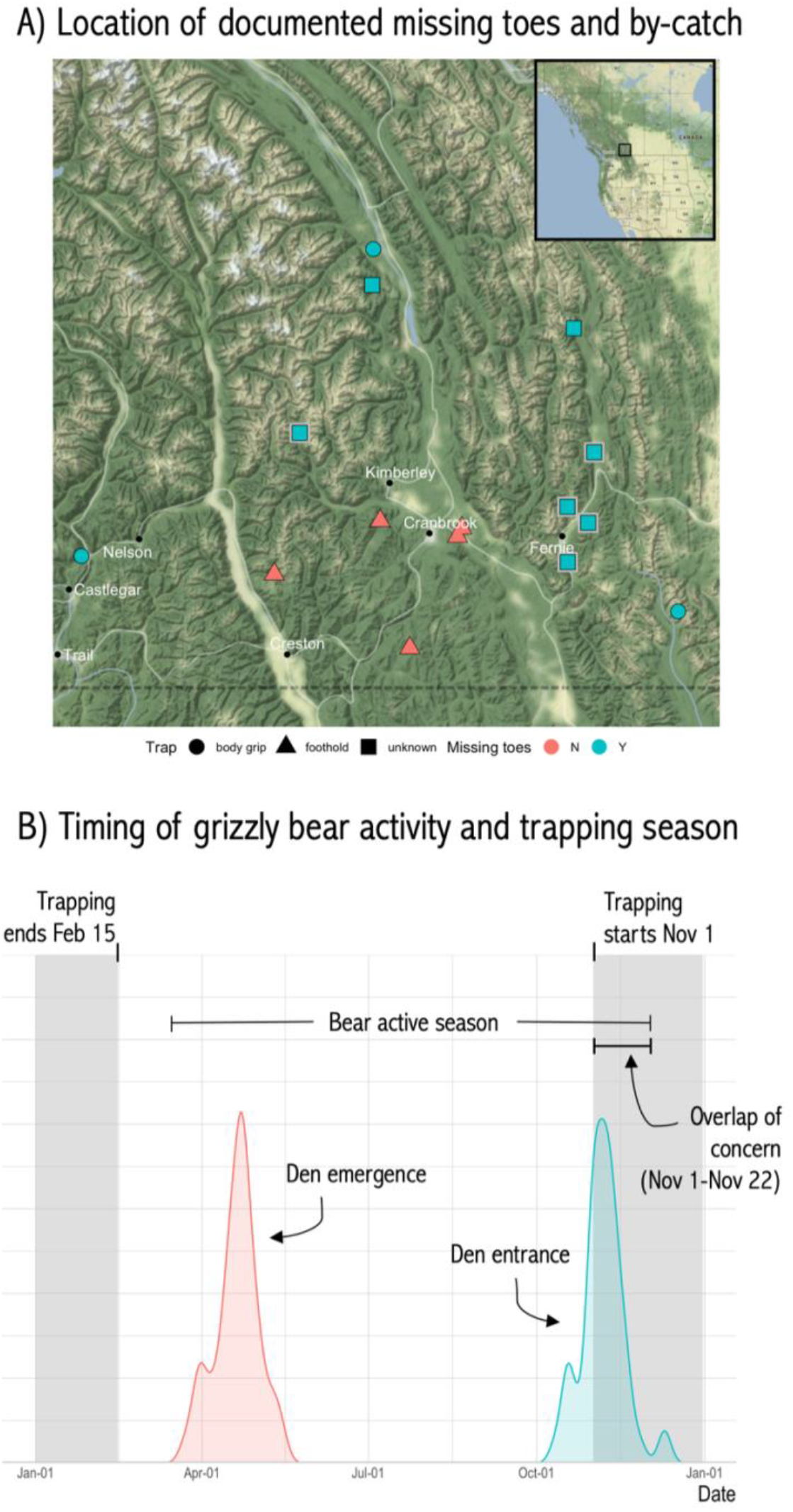
A) Locations of observations of bears with amputated toes and bears with traps on their feet. Detections made through research are outlined in grey, while detections opportunistically gathered throughout southeast British Columbia are not outlined in grey. B) The grizzly bear active season, defined from 61 collared bear den emergence and entrance dates, with the marten trapping season (November 1 to February 15) shown in grey. The 95^th^ percentile of den entrance dates was November 22. We suggest that December 1 represents an evidence-based start date for trapping to reduce conflicts between bears and trappers.

Like humans, grizzly bears have twenty digits. Of the 59 individual bears captured, 4 (~7%) were missing some of the toes on one of their front feet (Fig 2). In all cases, the toes lost were adjacent and terminated in a linear plane, and the injury had healed. At the time of their capture, these animals were all between the ages of 4 and 12 years old. Three of the four captured bears with amputated toes were male, while the larger collar sample had slightly more females (n=31) than males (n=28). The female with amputated toes was captured with a yearling cub, suggesting that amputated toes did not preclude reproduction. Based on radio collar telemetry data, these animals moved in a similar fashion to bears with all their digits. While toe loss did not appear to impact their locomotion, it may have influenced conflict behaviour, which was common in these animals; three of the four animals with amputated toes were involved in human-bear conflicts. One of these four bears was killed shortly after capture while breaking into a rancher’s calf pen, another was likely involved in an attack on a human, and a third was captured by conservation officers in late fall after complaints of bear issues on a farm. While the sample size for this conflict behaviour is small, less than one third of the bears with all their toes (n=55) died or were involved in conflict during the time they were collared. The reason for elevated conflict behaviour observed in the bears with amputated toes could be related to challenges feeding on natural foods at certain times of year, such as digging roots during spring and fall (McLellan and Hovey 1995), or these bears could be naturally bolder individuals and thus more likely to become both trapped and involved in future conflict. Alternatively, the pattern of increased conflict could be an artefact of the small sample sizes we had available (n=4). Nevertheless, while we had identified a concerning trend of amputated bear toes but the healed wounds left us little evidence for the cause of the toe loss.

**Figure 2.**
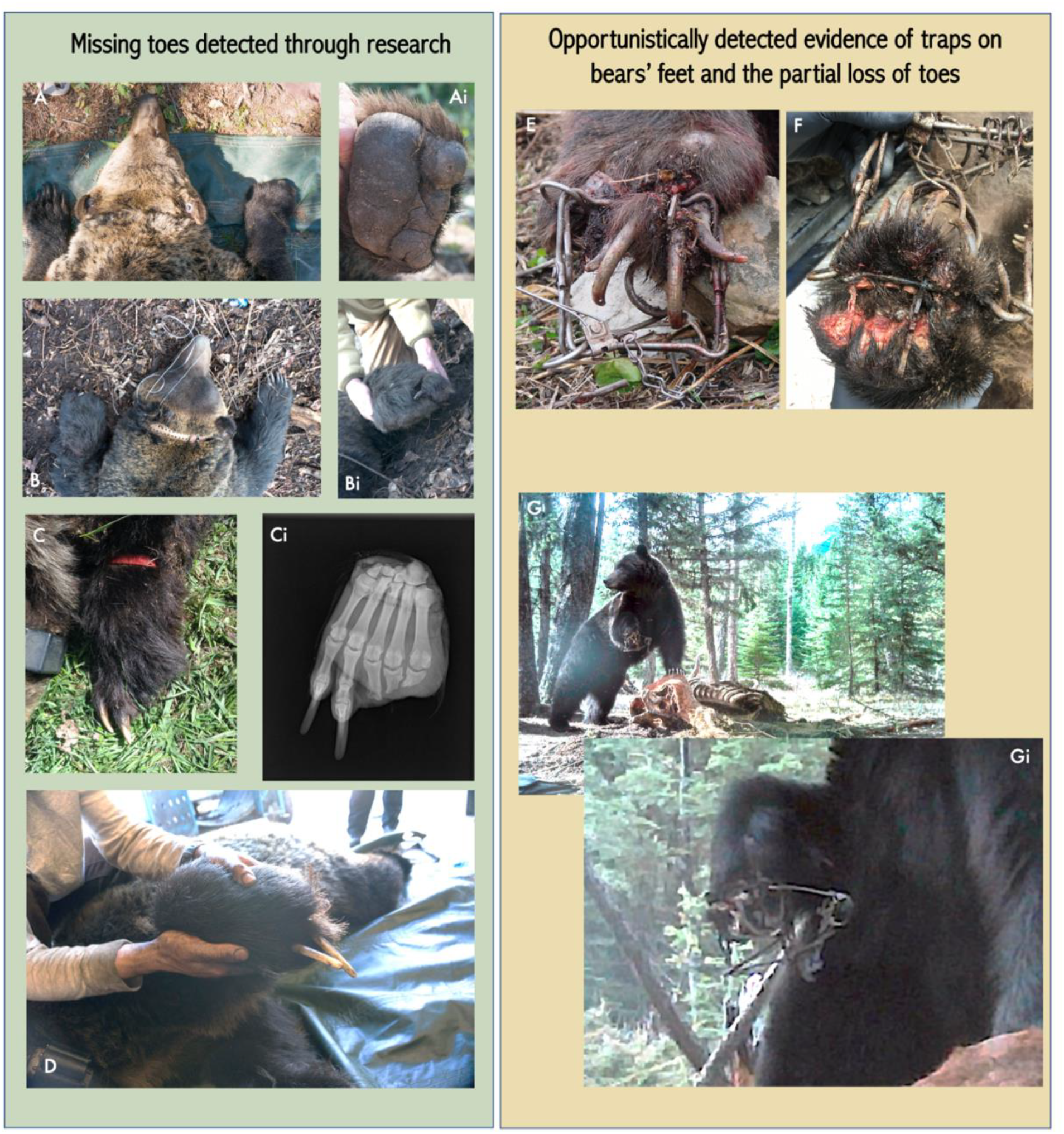
Animals detected with amputated toes through research captures in the Elk Valley (A-D), and opportunistic evidence of grizzly bears with traps on their feet (E-G). Each letter in A-D represents one of the four unique individuals detected with amputated toes. Ci) an X-ray of the foot of EVGM55, who was killed in a conflict with a cattle rancher. E) a body grip trap on the foot of a hunter-killed grizzly bear from the Flathead Valley. F) the paw of a 1-year-old grizzly bear stuck in a body grip trap captured by conservation officers near Pass Creek due to conflict behaviour from a mom and offspring; significant infection was present, and the toes had nearly completely separated from the foot by the time of capture. G) a grizzly bear with a body grip trap on its foot detected feeding on a cow carcass by a remote camera west of Invermere.

## SEARCHING FOR CAUSE

We first consulted with veterinary professionals to determine if the amputated toes could be congenital (i.e., lost before birth and due to natural causes). After review of multiple photos and X-rays (Fig 2) there was little evidence to suggest that this was congenital. On the contrary, strong evidence for injury post-birth included clear evidence of toe bone fracture and healing, and consistent linear wounds suggesting amputation related to human sources. As a result, we proceeded to search for a human-caused source of the issue.

Southern British Columbia is fortunate to have a long history of people interacting with grizzly bears through research, human-wildlife conflict work, hunting, guide outfitting, trapping, and simply living in bear country. After identifying the amputated toe issue, we spoke with First Nations, scientists, conservation officers, wildlife managers, guide outfitters, and trappers, all with lifetimes of experience on this land. A common topic in these discussions was grizzly bears being incidentally caught in baited traps set for furbearers, particularly in traps set for marten (*Martes americana*) and wolves (*Canis lupus*). Indeed, non-target captures stemming from legal trapping activities have been reported for many species. Cougars (*Puma concolor*) have been incidentally captured in foothold traps or snares set for wolves and bobcat (*Lynx rufus*) (Knopff et al. 2010, Andreasen et al. 2018), and free-ranging cats and dogs and even birds have been incidentally caught in furbearer traps (White et al. 2021). We knew that grizzly bears could be caught in foothold traps set for wolves, given that in recent years several bears had either been killed in, or required release from, wolf traps in southern BC; however, we were less familiar with the role that body grip traps could play in the loss of toes. Body grip traps are killing traps designed for furbearers such as marten, skunk (*Mephitis mephitis*), fisher (*Pekania pennanti*), weasel (*Mustela sp.),* muskrat (*Ondatra zibethicus*), beaver (*Castor canadensis*), bobcat, and lynx (*Lynx canadensis*) (Fig 2). Trapping legislation in BC allows for the use of a variety of trap models made by different manufacturers, provided the devices reliably and humanely kill their target species (Province of British Columbia 2021, White et al. 2021). In our region of interest (Fig 1), small body grip traps (commonly referred to as 120 conibear traps) are most often used to capture marten and weasels, and they are typically set at the mouth of a baited wooden box that is affixed to a tree at a height of approximately 1.5 meters from the ground.

In the spring of 2018, we deployed four body grip traps, complete with their boxes and bait (usually a small piece of meat), on trees to see if bears would investigate these traps. We wired the traps in such a way that they could fire but not close and trap the bears. We monitored the traps with remote cameras for approximately two weeks to see if bears would visit and spring the traps. Grizzly bears visited all four traps and sprung two of them. Pictures and videos showed bears investigating the traps and manipulating the boxes with their paws (Fig 3). Video showed that one of the traps was set off with a bear’s nose, but the cause of the other trap being set off could not be determined because we only had pictures, not video, of bears investigating the trap. Although this was a small trial with low sample sizes, younger bears seemed to investigate and set off traps more often than older bears. Even with the small sample, it was clear that baited traps attracted bears and that bears set off the traps as they tried to get the bait. We heard from some trappers that they voluntarily delayed the start of their marten trapping season (which legally opens November 1, Province of British Columbia 2021) to avoid having bears wreck their trap sets, further confirming that bears are attracted to these common sets.

**Figure 3.**
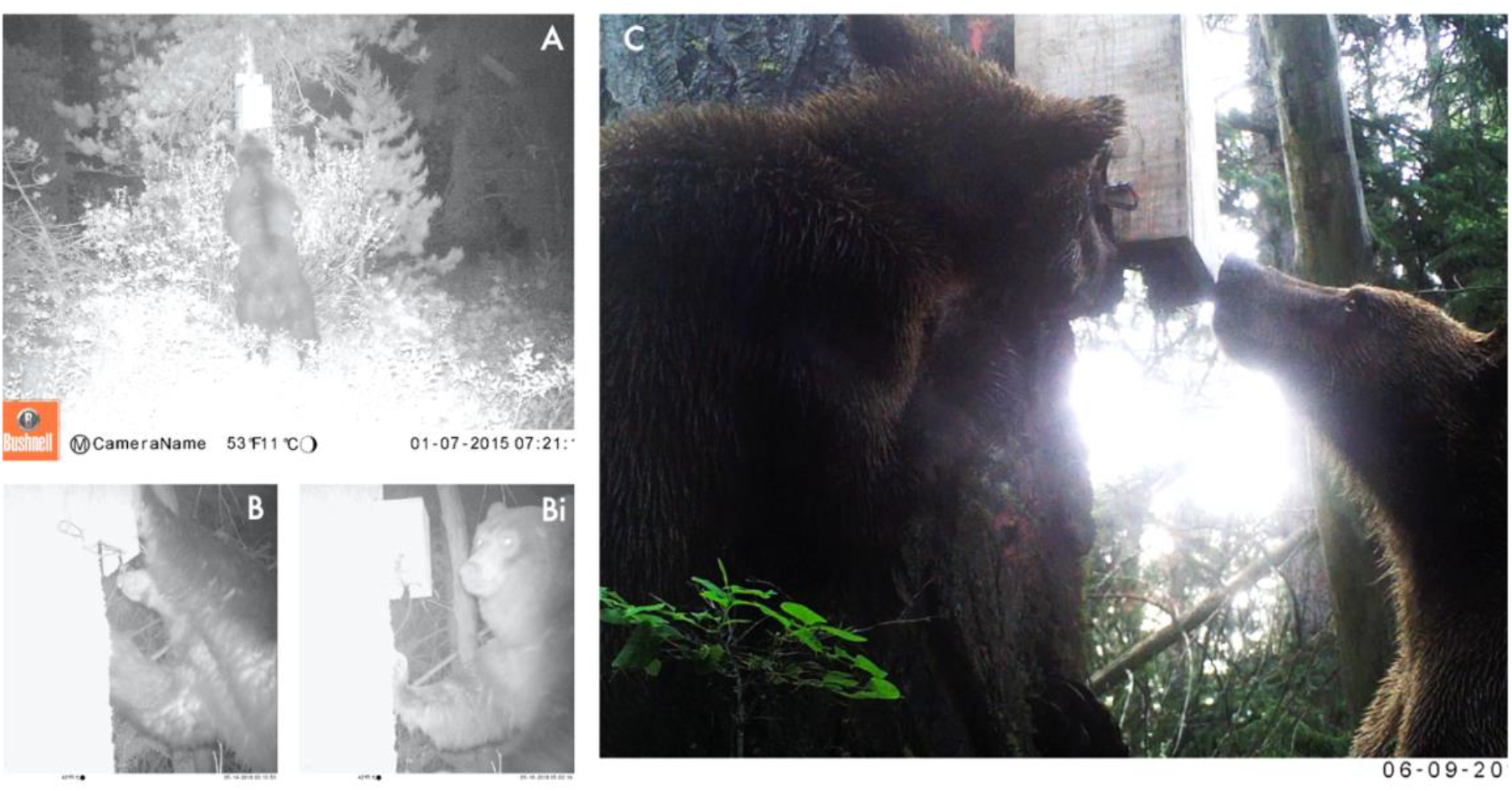
Grizzly bears detected at trial marten body grip traps. The traps were wired so they could spring but not close on bears’ feet, and they were set for two weeks in areas frequented by bears. Investigatory behaviour was common, and two of the four deployments were set off.

We became more convinced that furbearer traps were the cause of the amputated toes as we accumulated anecdotes and photo evidence of grizzly bear feet stuck in traps (Fig 2). Photos came from hunters, remote cameras on ranches, and conservation officers, and they all suggested the same mechanism. A body grip trap was affixed just behind the toes and amputation was underway. Beyond identifying a likely cause for the amputated toes, the photo evidence and information from people out on the land revealed that bears with traps stuck on their feet and amputated toes was an issue beyond the extent of our study area. In addition to the records of body grip traps on bears’ feet in southeast BC (also known as the “Kootenays”), photos of a grizzly bear with a body grip trap on its foot were reported in Wyoming, USA in 2017 (McKim 2017), and brown bears with body grip traps on their feet are observed almost every year in Finland (Kai-Eerik Nyholm pers. comm.).

We were also aware of multiple reports of grizzly bears being caught in foothold traps set for wolves, and we believe this is another possible source of toe loss. Between 2010 and 2020, at least five grizzly bears in southern BC were caught in wolf foothold traps (with the trap often closing right behind the toes) and had to be released by conservation officers and scientists. However, in the records we were able to collect, body grip traps were the most common trap on bears’ feet where it was clear the trap was the cause of toe loss. In one case, it is suspected that a bear’s foot became stuck in a body grip trap set on the ground for skunks on private property. However, given the near ubiquity of marten trapping with box traps across our area (an average of 2,082 marten were harvested annually in the Kootenays from 2013-2017), and the broad occurrence of amputated toes and traps on feet, we focus on marten and weasel trapping as likely sources of most of the body grip traps that end up on bear feet.

We compiled data from collaring projects in adjacent areas within British Columbia to assess if a broader pattern of toe loss could be revealed. Collaring projects in the Selkirk and Purcell Mountains (near Nelson and Creston, Fig 1A) between 2004 and 2018, and in the Flathead Valley (southeast of Fernie, Fig 1A) between 1978 and 2020, captured 72 and 206 grizzly bears respectively. Of the 20 bears captured in the Purcell Mountains, one adult male bear (5%) fit the pattern of having well-healed but amputated toes, while none of the 52 bears captured in the Selkirk Mountains had amputated toes. However, grizzly bears were accidently captured by trappers in foothold traps set for wolves on at least three occasions during the study in the Selkirk and Purcell Mountains, but no evidence of toe loss due to incidental grizzly bear capture in footholds was reported. Likely because bears were either released from the traps or killed. None of the bears captured in the Flathead Valley had amputated toes; however, a hunter killed a bear just north of the Flathead study area with a body grip trap on its foot (Fig 2E). Results from these adjacent studies did not reveal any general insights into the observed spatial distribution of bears with amputated toes. This is not surprising given that traplines in British Columbia are distinct areas trapped exclusively by registered trappers, and variation in trapping intensity, target species, and timing creates variable exposure of bears to body grip traps across the landscape. Additionally, unsecured attractants such as apple trees, garbage, roadkill, and livestock may increase bear use of low elevation areas and delay den entry, perhaps increasing the chance of overlap with trapping activities in these areas.

Observations of bears stuck in body grip traps and bears with amputated toes have mainly been documented in the past 15 years and are not restricted to a single area or trapline, but rather occur broadly at low frequency across the Kootenays and beyond. This increased detection of bear toe loss could be related to an increase in grizzly bear research and monitoring; however, opportunities to identify these issues have been abundant for several decades as many bears were captured by conservation officers, harvested by hunters, and monitored for research purposes in the Flathead. It is possible that in recent years the increased use of stronger traps, as mandated by evolving humane trapping standards, has created a situation where bears now have more difficulty extracting themselves from body grip traps when captured.

We summarized the den entry and exit dates for 61 animal-years in the Elk Valley to assess when bears were denning in the fall. We found the median den entry date (i.e. the date when 50% of bears had denned) was November 6, and the 95^th^ quantile of den entry was November 22, revealing a critical 3-week period of overlap between active bears and the trapping season. We found no overlap between den emergence and the end of the trapping season; thus, we focus on fall trapping as the source of the issue. More support for the idea that the concurrence of the fall active season for bears and the beginning of the trapping season is the main issue is that we have not seen similar problems of amputated toes in black bears, who den several weeks earlier than grizzly bears and would thus be less exposed to traps.

## HONING A SOLUTION

Most trapping of furbearers is done in winter when fur is prime (i.e., underfur is dense and guard hairs are long) and most valuable. Given that grizzly bears hibernate for the winter, trappers should generally be able to avoid accidently catching bears. However, some trapping seasons open in late fall, such as the marten season in much of British Columbia which begins November 1 and extends to February 15. Reducing the overlap between the period when bears are active and the trapping season is open is one way to minimize the amputation of bear toes and prevent trappers from losing their traps and having their sets destroyed by bears. Shifting the start of most trapping that coincides with the active bear season from November 1 to December 1 would essentially eliminate overlap between trapping and bears. This solution has been previously used in southeast British Columbia to avoid catching and killing bears in neck snares set for wolves, an issue first documented in the Flathead project (McLellan 2015, Province of British Columbia 2021).

Indeed, such a change to the rapping regulations to address the amputated toe issue was proposed in 2019 but was not implemented. Much of the trapped landscape in the Kootenays is rugged, mountainous terrain, and many trappers, especially in the west Kootenays, reported that they were effectively unable to access their traplines for much of the winter beginning in early December due to high avalanche risk and associated safety concerns. The trappers stated they did most of their trapping in November and early December, and that shortening the season by delaying the opening date to December 1 would largely remove their ability to harvest marten from their trapline. However, this was not a ubiquitous issue because many trappers in the east Kootenay reported voluntarily delaying their trapping until December to avoid bears, and other users of the landscape such as backcountry skiers and snowmobilers recreate in the mountains throughout the winter. Nevertheless, provincial biologists instead implemented a condition on all active trapping licenses in the Kootenay region stipulating that, beginning in 2021, all body grip traps set for marten during the month of November must be enclosed in a box with an opening no larger than 3 1/2 inches. This constricted entrance was thought to be narrower than most bear paws. The license condition also recommended that, prior to December 1, trappers should use these same boxes to enclose similarly sized killing traps set for other species on dry ground. The efficacy of the modified enclosure entrance in eliminating bear toe loss has not yet been tested, but these modified marten trap boxes are believed to be sufficient to ensure grizzly bears are not able to access a set trap. Because there is more uncertainty in the effectiveness of the modified enclosure approach than there is in delaying the start of the trapping season, we recommend that compliance and efficacy is monitored to ensure this intervention is effective and uptake is high. Such monitoring would allow for adaptive management and changes to the approach as needed to ensure a successful outcome for bears and trappers alike.

Additional approaches have been suggested to help resolve the issue, but we do not recommend their implementation. For example, one option is to sufficiently anchor traps so a bear that is incidentally caught can hopefully pull its foot free. To address the viability of this solution we conducted a small experiment using front appendages of deceased bears and four types of traps approved for marten trapping in British Columbia (Province of British Columbia 2021). Our experiment consisted of three phases: the first used before and after X-rays to assess whether traps could immediately break bear toe bones, the second assessed whether bears could easily pull their feet free from traps (which we simulated by solidly anchoring the trap and having an 86 kg person attempt to free the feet by pulling on the leg with maximum human pull), and the third assessed how much pull (kg) was required to free the feet. We gathered the front appendages from one adult and two cub grizzly bears killed in the Elk Valley during fall 2021 and tested four approved body grip traps: 1) Bélisle Super X 120, 2) Northwoods 155, 3) Sauvageau 2001-5, and 4) Sauvageau 2001-6.

Results from our experiment suggested that traps do not immediately break bones, but bears were unable to easily extract themselves from these traps. X-ray results confirmed that setting traps off on adult or cub feet did not break or fracture any bones. Despite using maximum human effort and both sustained and jerking pulls, we were only able to free adult grizzly feet from traps 20% (95% CI: 7-33%) of the time, and cub feet 63% (95% CI: 53-72%) of the time. For 80% of our attempts with adult feet and 37% of our attempts with cub feet, we were unable to free the feet from the traps despite repeatedly jerking with as much force as we could muster. Considerable variation in release efficacy was observed between trap models (Table 1), where the Sauvageau traps generally released bears the least frequently, and Northwoods 155 released bears the most. Finally, we used a come-along or a truck, with a scale affixed inline, to determine the steady pull required to extract an adult foot from the Sauvageau traps. The average pull required to free a foot was 164 kg (range=91-232 kg). Even after maximum pull was applied, we did not detect superficial damage to toe bones or joints, although clearly significant pain would be sustained by the bears during this time, and we are not certain that bears could always muster the >230 kg pull required to ensure reliable release.

**Table 1.** Phase two results of the toe extraction experiment where maximum human pulling ability was applied in an attempt to free grizzly bear feet from traps.

| Age | Trap | Trials | Percent released |
| --- | --- | --- | --- |
| Adult | Bel120 | 5 | 0 |
| Adult | North155 | 5 | 60 |
| Adult | Sauv160 | 5 | 0 |
| Cub | Bel120 | 5 | 100 |
| Cub | North155 | 5 | 100 |
| Cub | Sauv120 | 8 | 50 |
| Cub | Sauv160 | 6 | 0 |

**Table 2.**
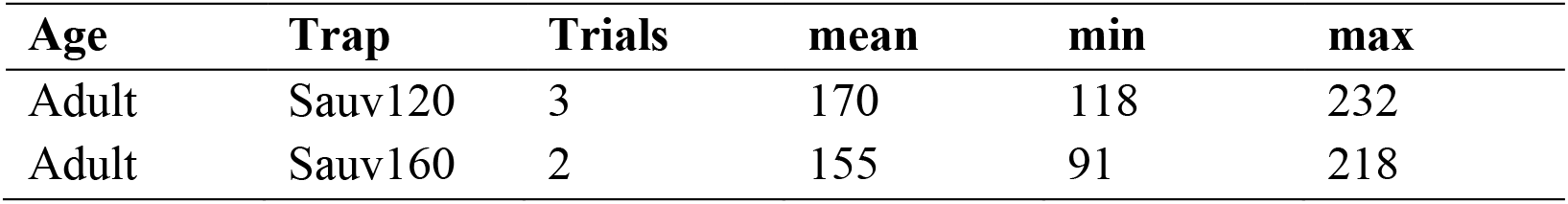
Phase three results of the toe extraction experiment where a steady pull from either a come along or a truck was applied to extract feet from traps. The maximum pull (kg) recorded on an inline scale prior to release was recorded for each trial.

| Age | Trap | Trials | mean | min | max |
| --- | --- | --- | --- | --- | --- |
| Adult | Sauv120 | 3 | 170 | 118 | 232 |
| Adult | Sauv160 | 2 | 155 | 91 | 218 |

Based on the results of our experiment, we believe that better anchoring traps would present safety concerns for bears, trappers, and wildlife professionals that might be called to release bears. Although there is little work assessing the pull of a bear, a strong animal such as a horse provides some insight into the forces animals can generate. A horse can exert ~70% of its body weight at max exertion (Smith 1896). The adult female grizzly bear whose appendages we used weighed approximately 140 kg, which would equate to a dead pull of ~98 kg if we use the weight to exertion relationship from horses. This would be well below the maximum of >230 kg required to reliably extract feet from these traps, and barely above the minimum recorded of 91 kg. As a result, bears may often be unable to escape from well anchored traps. There is a greater chance that bears could free themselves if a long leash (>2 m) is used so bears could run and generate more pull. It is thought that a 140 kg bear could generate between 400-1000 kg of pull with a running start (Flaa et al. 2009), however, significant damage to bears’ feet and traps is possible in this case and we do not recommend this solution. Overall, our experiment suggests that bears cannot easily extract their feet from traps and better anchoring traps presents significant risk to people and bears. Furthermore, current regulations for trap check intervals on killing traps such as body grips can be as long as 14 days in British Columbia and would require significant reduction (<24 hours) if bears could become permanently held in traps. Such a change in trap check intervals from 14 days to 24 hours would render most large winter traplines impractical for trappers.

The most viable solution to the amputated toe issue requires that bears’ feet do not enter traps at all. Changing the season length is the most reliable solution, but possibly reduces trapper access to trapping opportunities in portions of the Kootenays. Constricting the entrance size of the trap presents a possible simple solution but requires further testing to ensure the design is effective at excluding bears while not dissuading marten from entering. Modified entrance size requirements will also require compliance monitoring, adding additional responsibilities to already overtasked Conservation Officers. The solutions we present here have various pros and cons, and we hope that this work can help policy makers make an informed decision and choose a solution that will resolve the amputated toe issue while ensuring trappers have sufficient opportunity to trap furbearers.

Trappers have a long history of supporting conservation efforts and are sentinels of changing landscapes and wildlife populations. However, trapping of furbearers in British Columbia has received increased scrutiny amidst shifting sociopolitical trends in society. Bold and novel strategies will be required from trapping organizations and wildlife management agencies to maintain public support while also recruiting and retaining the next generation of trappers. A tactful strategy to uphold public support is to proactively assess and react to emerging evidence that threatens social acceptance of trapping (e.g., grizzly bears with amputated toes), and where changes could help increase the well-being of animals. This approach is the backbone of the North American Model, and while emerging evidence or science can sometimes conclude no direct threat to wildlife populations, organizations must equally weigh social ramifications and understand that the future of trapping may be predicated more on social support than consensus over sustainability. Here we document a troubling issue where grizzly bears are losing their toes after becoming stuck in baited furbearer traps. The timing of furbearer seasons is the primary issue, with trapping seasons opening weeks before all bears have denned, which creates a problematic period of overlap between active bears and baited traps. We provide evidence of the issue, solutions to remedy it, and ultimately urge immediate action.

## DATA AVAILABILITY

All analyses and supporting data can be found at https://github.com/ctlamb/Grizzly-MissingToes

## ACKNOWLEDGEMENTS

This work was conducted within ?amak?is Ktunaxa, the homelands of Ktunaxa people. We thank the Ktunaxa Nation for their support of our kǂawǂa (grizzly bear) work. Thanks to the trappers, guide outfitters, hunters, First Nations, and scientists that discussed this work with us, helped us identify the issue, and explore solutions. A special thanks to trapper Blair Chatterson who lent us traps, trapping supplies, and discussed this issue with us. Thanks to veterinarian Steven Chapman for X-rays of EVGM55’s foot, interpretation of these images, and discussion on potential causes. Thanks to veterinarian Cali Lewis and Tanglefoot Veterinary Services for before and after X-rays of bear feet for our experiment, interpretation of images, and many thoughtful discussions on the causes of, and potential solutions to, this issue. Thanks to Emily Chow and Stewart Clow for help setting traps and staunch efforts to free bear toes from these traps during the experiment. Thanks to Patrick Stent for support of our research project and many helpful discussions as we worked towards a solution to this issue. Thanks to Adam Ford and Melanie Dickie for suggestions on displaying results. Thanks to Garth Mowat, Mateen Hessami, and Tim Killey for suggestions on improving the manuscript. These data were collected with support from the Government of British Columbia, Habitat Conservation Trust Foundation, Vanier Canada Graduate Scholarship, Liber Ero Fellowship Program, National Science Engineering and Research Council, Fish and Wildlife Compensation Program, Forest Enhancement Society of British Columbia, Teck Coal, Columbia Basin Trust, Counter Assault, Wildsight, Nature Conservancy of Canada, Wildlife Conservation Society, Yellowstone to Yukon Conservation Initiative, Safari Club International, Sparwood and District Fish and Wildlife Association, Elkford Rod and Gun Club, Ministry of Transport and Infrastructure, Outdoor Research, Patagonia, and the British Columbia Conservation Officer Service.

